# A large 3D-printed integrated lens-biprism element enhances contrast in transmission stereomicroscopy

**DOI:** 10.1101/2025.05.20.655140

**Authors:** Liam M. Rooney, William B. Amos, Shannan Foylan, Jay Christopher, Charlie Butterworth, Ralf Bauer, Gwyn W. Gould, Gail McConnell

## Abstract

Stereomicroscopes are routinely used across disciplines for material and surface characterisation due to their simplicity of use and minimal specimen preparation requirements. However, the stereomicroscope transillumination design is suboptimal, as a single incident beam at the specimen plane is shared and transmitted via two laterally offset detection axes. This flaw limits life science applications due to the transparent nature of samples which results in poor contrast images. We use a single additional element in the illumination path to correct the illumination uniformity across the field of view and, by doing so, enhance image contrast and facilitate detection of refractive structures in transparent biological specimens. We designed and fabricated an integrated lens-biprism element using low-cost, consumer-grade 3D printing methods and consumables. This 3D printed lens-biprism distributed diverging rays from a single incandescent light source into two parallel beams that converged at the specimen plane and transmitted through the respective left and right detection axes of a stereomicroscope. This improved transillumination setup increased the image contrast by up to 67.62% compared to the conventional stereomicroscope setup. We demonstrated the benefit of the lens-biprism element by visualising dynamic cellular events in live tissue and discerning refractive structures more easily in transparent specimens.

## Introduction

Stereomicroscopes are commonplace in many research laboratories, clinics and manufacturing facilities. The stereoscopic imaging approach provides a three-dimensional (3D) visualisation of multi-millimeter-sized specimens with little need for user alignment or arduous specimen preparation. Moreover, they do not invert the image, which is common among other optical imaging modalities. Traditionally, stereomicroscopes were produced in large quantities for use in the factory assembly of electronic devices but also see applications in metallurgy^1^ and mineralogy^2^, fractography^3^, forensic analyses^4^, and microbial phenotyping^5^. Stereomicroscopes are routinely configured per the Greenough-type setup, where two identical and symmetrical spatially offset detection paths are housed within the body of the microscope, each composed of independent objective lenses, zoom optics, prisms, and ancillary optics. The common main objective (CMO)-type is also routinely used, where light is collected by a shared objective lens and the image is formed through two laterally offset parallel optical paths containing zoom lenses, tube lenses, ancillary eyepieces and/or prisms to direct light to a camera detector. Other stereomicroscope designs, such as those using telecentric lens systems, are less common^6^. The Greenough stereomicroscope facilities high numerical aperture, ergo higher spatial resolution but, due to the geometry of the offset detection axes, they suffer from image distortions owing to the non-paraxial optical arrangement^7^; this, combined with the inefficient transillumination regime of the stereoscope only guiding a fraction of the illumination light to the eyepieces presents an opportunity to adjust the microscope setup to improve image quality.

While rivalling the products of long-established microscope manufacturers in lens performance, stereomicroscopes were designed for epi-illumination and therefore are often poorly adapted for transillumination. This problem severely limits the application of these readily available and cheap microscopes in biomedical applications. The illuminator routinely comprises a single incandescent bulb placed beneath the specimen and separated from it by a diffusing screen. Thus, a single light source must serve the two optical paths on the left and right of the stereomicroscope. This arrangement has two issues; namely that the source position appears different in the two eyepieces because of the parallax effect, and that the field is often incompletely illuminated.

The inhomogeneity of illumination over the field of view hinders the visualisation of refractive structures in thin or transparent specimens, as is routine in biological applications. Efforts to adapt stereomicroscope transillumination have been employed to improve image quality, but these typically require expensive instrumentation or specialist training. For example, previous methods include the use of fibre optics to improve stereo contrast^8^, optofluidic microlens arrays to increase spatial resolution^9^, or metamaterials to enhance the optical performance of the stereomicroscope^10^. We propose the introduction of a low-cost integrated lens-biprism element into the transillumination path of the stereomicroscope to provide converging collimated illumination, thus improving the homogeneity of illumination over the field and enhancing contrast to discern biological structures. This custom optical element effectively combined a large convex lens to collect and collimate diverging rays from the in-built light source, and a biprism with a refractive angle sufficient for two converging beams to intersect at the specimen plane. This made for a disk with flat prismatic surfaces on one side and a convex spherical surface giving focusing power on the other. Due to the bespoke form and prescription of this large optical element, we used previously validated, low-cost, consumer-grade 3D printing and post-processing to fabricate the lens-biprism element, which has recently been employed to fabricate high-quality 3D printed optics for various optical imaging applications including multi-element objectives^9^ and in entirely 3D printed microscope systems^11^, and have been rigorously validated using classical optical methods^12^. We evaluated our approach using a conventional Greenough-type stereomicroscope and demonstrated the improvement in illumination and contrast using transparent refractive structures.

## Materials & Methods

### Background Theory for Enhanced Contrast using an Integrated Lens-Biprism Stereomicroscope Transilluminator

Biological specimens often exhibit local variations in refractive index *n*(*x*,*y*), which cause small angular deflections in transmitted rays as they traverse regions with non-uniform optical properties. These angular deviations are the physical basis for contrast in label-free imaging of transparent samples. In geometrical optics, the deflection of a ray in a medium with a spatially varying refractive index can be described by a simplified scalar form derived from the eikonal equation^13^. For a ray propagating predominantly along the z-axis and encountering a transverse gradient in the x- direction, the angular deviation θ(*x*) from the original propagation direction is given by Equation 1, where *n*_0_ is the ambient refractive index and 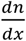 is the transverse refractive index gradient at position *x*. This approximation is valid in the small-angle (paraxial) limit and assumes weak phase gradients.

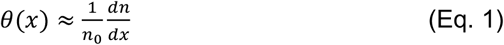

The enhanced illumination provided by the biprism ensures that such angular deflections modulate the intensity of light entering the objective lens. A ray deflected by θ(*x*) will be laterally displaced at the image plane as descried by Equation 2, where *f* is the effective focal length of the imaging lens.

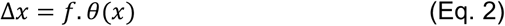

As stereomicroscope objectives have finite numerical apertures, rays that are deflected significantly from the optical axis may fall outside the collection cone or be redirected deeper into it, resulting in detectable changes in local image intensity.

Assuming the image intensity *I*(*x*) is proportional to the local density of rays arriving at the detector, and that angular deflections vary smoothly across space, the intensity can be modelled by Equation 3.

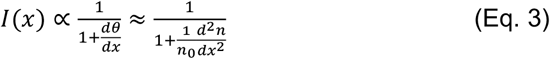

Equation 3 shows that image contrast is directly related to the second derivative of the refractive index. Structures with high curvature or sharp transitions in *n*(*x*) produce more significant angular spreading or convergence of rays, translating into increased brightness or shadowing. Since stereomicroscope illumination is incoherent, contrast enhancement could arise entirely from controlled angular light redirection, not interference. We propose that introducing a tailored angled illumination regime would enable higher-contrast imaging of refractive structures by transforming a change in refractive index into observable intensity variation.

### Design of an Integrated Lens-Biprism Element for Stereomicroscope Transillumination

The lens-biprism element was designed as a single integrated optical component comprising a convex first surface coupled to a circular-based biprism (Figure 1a). The second surface comprised two planar faces at the necessary wedge angle to split the incident beam into two converging parallel beams. These two parallel beams meet at the specimen plane and then diverge, with each transmitted beam propagating through further optical elements in the respective detection path of a stereomicroscope (Figure 1b).

**Figure 1.**
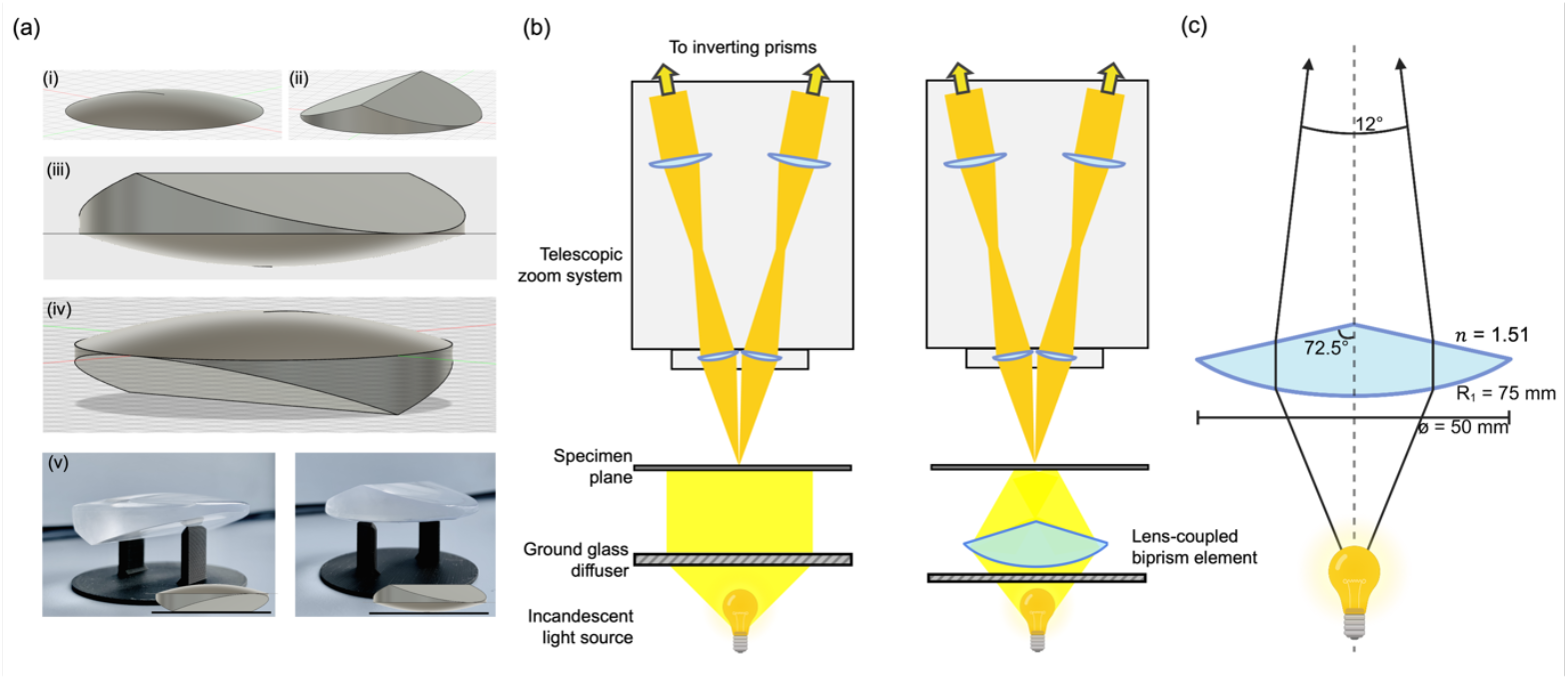
A large 3D-printed integrated lens-biprism element for improved stereomicroscope transillumination. **(a)** Renders of the 3D printed lens-biprism element showing the convex first surface (i), the biprism second surface (ii), the final coupled form in an upright (iii) and inverted (iv) orientation, and images of the 3D printed biprism element for the previous orientations (v). **(b)** A simplified schematic comparing the transmission paths of a conventional Greenough-type stereomicroscope with a standard transillumination setup and a biprism transilluminator. **(c)** A ray diagram illustrating the prescription of the lens-biprism element. Diagrams are not presented to scale.

The first surface was configured as a convex spherical surface designed to collimate the light radiating from a small source into an approximately parallel beam. The radius of curvature, *R*_1_, of the first surface was calculated using Equation 4 and the thickness, *T*, of the convex lens portion of the bulk element was calculated using Equation 5; where *n*_0_ = the refractive index of air, *n*_R_ = the refractive index of the 3D printed resin^12,14,15^, *f* = the focal length of the convex component, *R*_2_ = the radius of curvature of the second surface of the convex component (*R*_2_ = ∞ for planar surfaces, as would be the case here), and *D* = the diameter of the component.

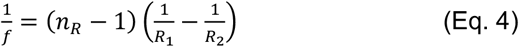

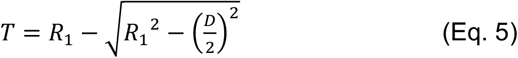

The lens-biprism element was designed to be placed in the transillumination path at 147 mm from the specimen plane, meaning *R*_1_ = 75 mm; thereby, *T* = 4.29 mm. Given the required focal length of 147 mm and the known refractive index of the 3D printing resin, *n*_R_ = 1.51, the wedge angle of 72.5° was calculated for the biprism component (Figure 1c).

### Fabrication of a 3D Printed Integrated Lens-Biprism Element

A 3D schematic of the biprism element was created using Fusion (v2.0.20970; Autodesk, USA) according to the geometry calculated using Equations 4 and 5, and fabricated using modified protocols from the ones described previously^11,12,14^. The polygon count was increased to the maximum available before exporting the model as a .STL file. The 3D print file was generated by importing the .STL file in LycheeSlicer (v5.2.201; Mango3D, France), encoding the print parameters, and exporting as a .CTB file. The biprism element was printed using a Mars 3 Pro 3D printer (ELEGOO, China) with photopolymerising clear resin (RS-F2-GPCL-04; Formlabs, USA) using an exposure setting of 9 seconds for a 10 μm layer height. The printed biprism element was post-processed by washing with neat isopropyl alcohol for 9 minutes, drying using an air duster (CA6-EU; Thorlabs, USA), and curing for 15 minutes using 385/405 nm light in a curing station (MercuryX; ELEGOO, China).

Custom spin chucks were designed to secure the biprism element during the post-processing steps required to render the optical faces smooth and transparent for imaging. A negative impression of the convex and biprism components was incorporated into two separate spin chucks designed to fit into a spin coater (Ossila, UK). The chucks were designed using Fusion, exported as a .STL file, and processed as a .3MF file for 3D printing using a fused filament fabrication (FFF)-style printer (P1S; Bambu Lab, China). The chucks were printed using Bambu PLA Matte filament using a 0.4 mm nozzle and standard system presets for printing with a 0.2 mm layer height and 15% ‘grid’ infill on a Smooth PEI build plate.

The convex first surface was processed first by carefully push-fitting to secure the element into the 3D-printed spin chuck. The surface was rendered smooth by spin coating a thin layer of liquid clear resin (version 4; VidaRosa, China) for 10 seconds at 2000 rpm, and post-curing for 10 minutes. The planar faces of the biprism component were then processed as above, but by depositing the thin layer of liquid resin along the length of the refracting edge of the biprism.

### Creating a Modified Biprism Transilluminator to Enhance Stereo-contrast

The final 3D printed lens-biprism element was incorporated into the transmission path of a trinocular Greenough-type stereomicroscope (XTL-3400; Shanghai Jinnshine Photonics Technology Co. Ltd., China). The integrated lens-biprism element was placed at approximately 75 mm from the incandescent light source, supported by a custom jig, and at 147 mm from the specimen plane. The stereomicroscope was coupled to a CMOS camera (Hypercam 183M; Altair Astro Ltd., UK) and images were acquired using Altair Capture (v. x64/4.7.14803.20190605, Altair Astro Ltd., UK). The camera was set at γ = 1 to preserve the same luminance to display brightness for all exposure settings.

### Assessing the Illumination Homogeneity of a Biprism Transilluminator

The image pairs were acquired from the left and right optical axes to determine the homogeneity of illumination. Briefly, a glass slide with a printed 2 mm x 2 mm grid featuring a single ink spot for reference was imaged using the stereomicroscope in the default setup. Image pairs were also acquired with three layers of 220 grit ground glass diffuser plates (45-655; Edmund Optics, USA) between the light source and the specimen. Image pairs were then acquired using three diffuser plates and with the integrated lens-biprism element in position as described above. Acquisition parameters were maintained for all conditions and images were obtained at the lowest zoom level of the microscope. Due to the design of the stereomicroscope, with the photoport fixed on the left-hand side of the eyepieces, the right-hand side images were acquired by rotating the whole microscope relative to its base by 180° while leaving the illumination assembly and sample fixed in place, and the images were digitally rotated by 180° to ensure that the two images were in the same orientation.

### Specimen Preparation and Imaging

A dissociated goldfish scale (*Carassius auratus*) and the abaxial epidermis of a white onion (*Allium cepa*) were selected to demonstrate the contrast improvement of the biprism transilluminator due to their fine structure, optical transparency, and difficulty to image using routine label-free imaging. The goldfish scale was removed and mounted in high-index resin (Fluoromount-G; ThermoFisher, USA)^16^, which also increased its transparency. The onion epidermis was prepared by mounting in phosphate-buffered saline (Gibco, pH 7.4; FisherScientific, USA) immediately before imaging. Images of the goldfish scale specimen were acquired as described above, with images of the whole scale acquired at a microscope zoom setting of 0.7× and a magnified region acquired at 1.2× zoom. Images of the onion epidermis were acquired at a microscope zoom factor of 4.5×. Time-lapse movies of the onion epidermis were acquired at a frame interval of 6 seconds using the acquisition settings described above and are presented in the supplementary materials.

### Image Processing and Analyses

All image processing and analyses were conducted using FIJI^17^ (v1.54f). Image pairs of the grid specimen were aligned and registered by drawing a linear region of interest (ROI) along the same grid line in each paired image and using the *Align Image by Line ROI* plugin in FIJI. A composite of each image pair was generated for the three transillumination setups described above, and the average intensity across the field of view was measured for both the left and right optical axes.

The contrast improvement was quantified by measuring the relative intensity of the growth annuli on the distal surface of the goldfish scale. A linear ROI (width = 5 pixels) was drawn parallel to one of the radial ridges on the scale, and an intensity plot profile was generated for both the triple diffuser and lens-biprism element transillumination setups. The contrast improvement was quantified using a custom Python script^18^ (Python v3.12) to measure the peak-to-trough value of the intensity peaks corresponding to the growth annuli.

It is important to note that the image data presented was not subject to contrast adjustment to demonstrate the improvement in contrast resulting from the lens-biprism transilluminator.

## Results

### A 3D Printed Lens-Biprism Transilluminator Improves Field Illumination

Figure 2 compares the illumination over the field-of-view (FoV) for a standard stereomicroscope setup and two modified setups: one with three layered ground glass diffusers and one with the 3D printed integrated lens-biprism element plus diffusers. We observed that the standard transillumination setup resulted in restricted illumination and parallax so that the source appears in stereovision far behind the specimen (Figure 2a, b). The addition of diffusing plates improved the spread of illumination, producing defined left and right-hand side intensity maxima, but did not entirely remove parallax (Figure 2c, d). In adding a integrated lens-biprism element to the illumination path, we observed superior coverage of the FoV with more uniform intensity and reduced parallax when compared to the previous setups (Figure 2e, f). The transilluminator setup incorporating the lens-biprism element provided illumination uniformity over a 15 mm FoV.

**Figure 2.**
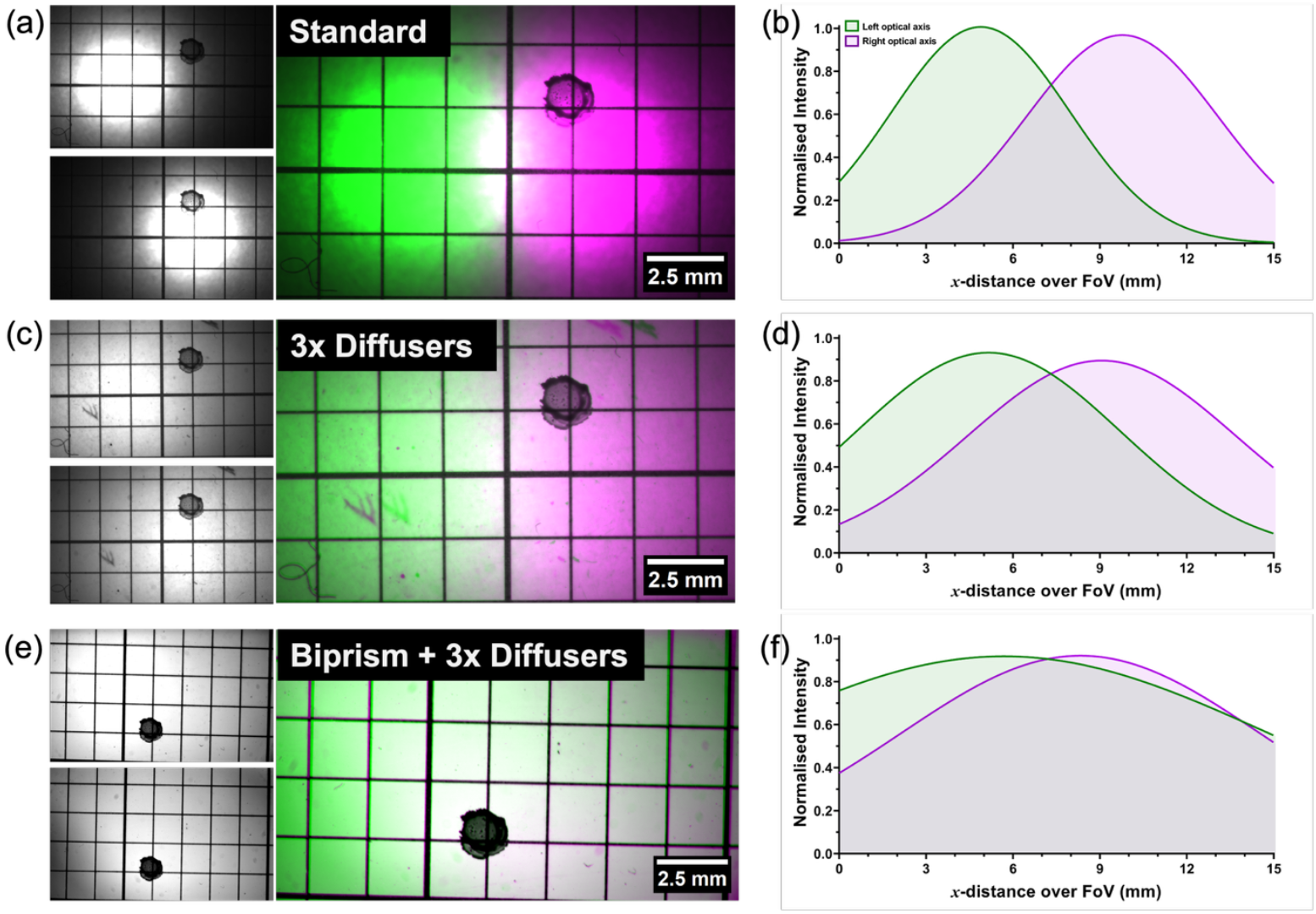
Improving the homogeneity of illumination of a stereomicroscope by using a 3D printed biprism transilluminator. **(a)** Images of a 2 mm grid marked with a single ink spot are presented with separate images of the left (top) and right (bottom) optical axes in a standard transillumination stereomicroscope setup. A false-coloured composite image of the left (green) and right (magenta) optical axes is presented, where ‘white’ signal corresponds to areas of homogeneous illumination. **(b)** A normalised intensity plot profile over the field-of-view shown in (a) fitted with a Gaussian function for the left (green) and right (magenta) optical axes of the standard stereoscope setup. **(c)** Images as presented in (a) for a transillumination setup featuring three diffusers placed in the illumination path. **(d)** A normalised intensity plot profile as described in (b) for a transillumination setup featuring three diffusers placed in the illumination path. **(e)** Images as presented in (a, c) for a transillumination setup featuring a 3D printed biprism element and three diffusers placed in the illumination path. **(f)** A normalised intensity plot profile as described in (b, d) for a transillumination setup featuring a 3D printed biprism element and three diffusers placed in the illumination path.

### Homogeneous Illumination Using a 3D Printed Biprism Transilluminator Enhances Contrast

Improving the uniformity of illumination over the FoV provided enhanced contrast, facilitating the visualisation of subcellular structures otherwise unobserved by conventional label-free stereomicroscopy. Figure 3 shows a goldfish scale imaged using a standard transillumination setup with three layered diffuser plates (Figure 3a) and with a biprism transilluminator added to this setup (Figure 3b). The growth annuli were visualised as stepped banding structures following the contours of the scale and were observed more clearly using the biprism setup. A direct comparison of the intensity along a linear ROI demonstrated the increase in contrast facilitated by integrating the lens-biprism element into the illumination path of the stereomicroscope. We measured a normalised contrast over the ROI of 0.1986 (standard deviation = 0.1485) for the standard stereomicroscope arrangement, which increased to 0.3329 (standard deviation = 0.2047) with the addition of the lens-biprism element, providing a 67.62% contrast improvement. We measured 54 peaks along the measurement ROI in the triple diffuser setup and 52 peaks in the biprism setup, with this small difference likely due to feature merging in the lower contrast data.

**Figure 3.**
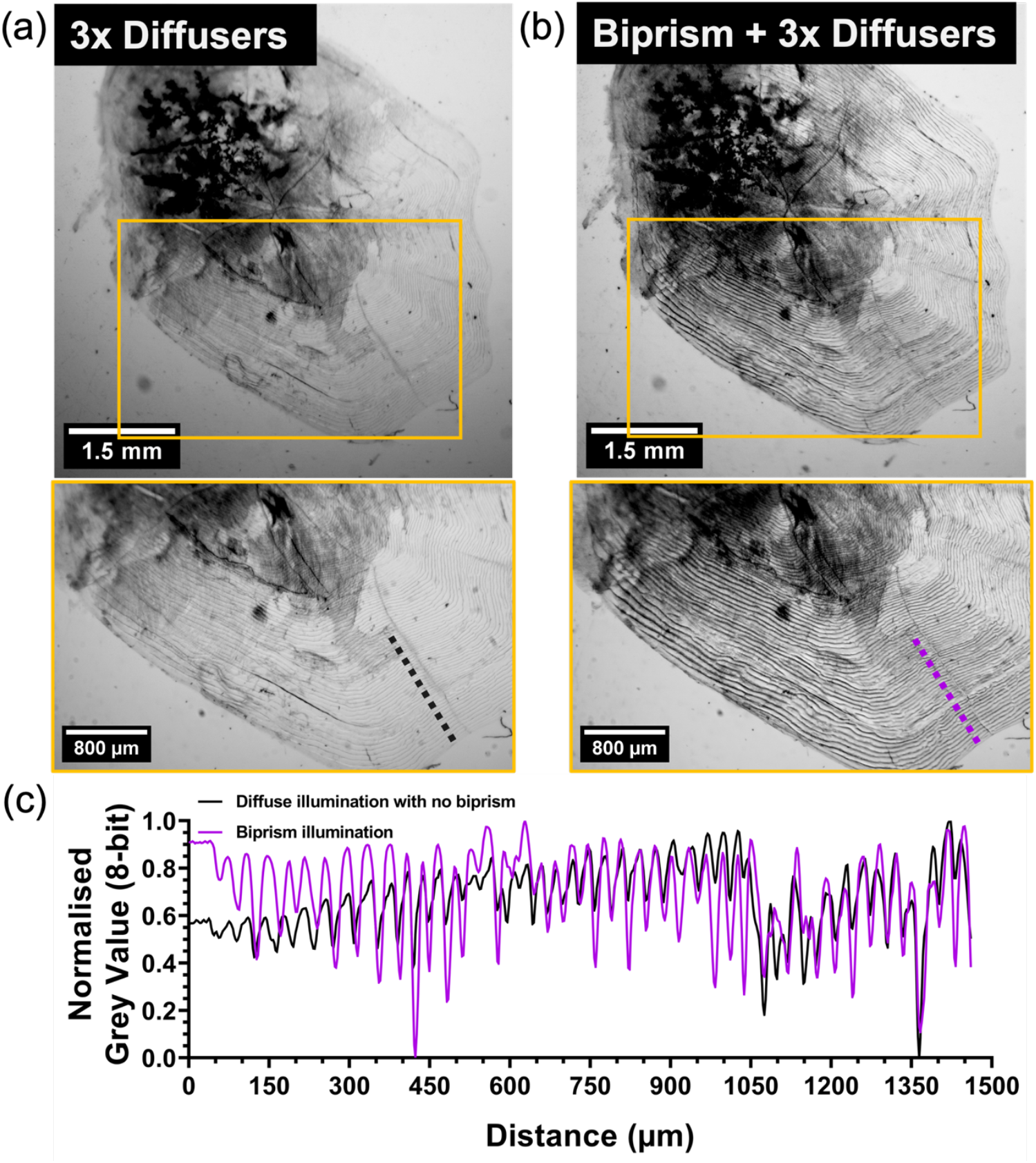
A 3D printed biprism transilluminator improves image contrast. **(a)** An image of a dissociated goldfish scale acquired using a transillumination setup with three layered ground glass diffusers. A magnified ROI is presented with a linear measurement ROI shown parallel to a radial ridge of the scale. **(b)** An image of a dissociated goldfish scale acquired using a 3D printed biprism transillumination setup. The same magnified ROI and measurement ROI as in (a) are presented. **(c)** A normalised intensity plot profile generated from the measurement ROIs presented in (a-b). The growth annuli are measured as dark bands along the radius of the scale. Black = diffuser setup, magenta = biprism setup.

We also imaged a live onion epidermis and compared the cellular structures and intracellular dynamics over time. We acquired time-lapse movies of the epidermis specimen using both the triple diffuser setup and the biprism transilluminator setup (Movie 1). The addition of the biprism element facilitated higher contrast, revealing dynamic behaviours that could not be easily observed by conventional stereomicroscopy such as cytoplasmic streaming, and movement of interstitial fluid through the apoplasm and middle lamellae. Movie 1 also compares a digitally magnified ROI showing these dynamics in closer detail. Figure 4a shows a single frame from Movie 1 acquired using a standard transillumination setup, and Figure 4b shows the same region acquired using a biprism transilluminator. The enhanced contrast facilitated visualisation of the nuclei across the field of view (green arrows) that were unobserved using a conventional illumination setup (Figure 4a, magenta arrows).

**Figure 4.**
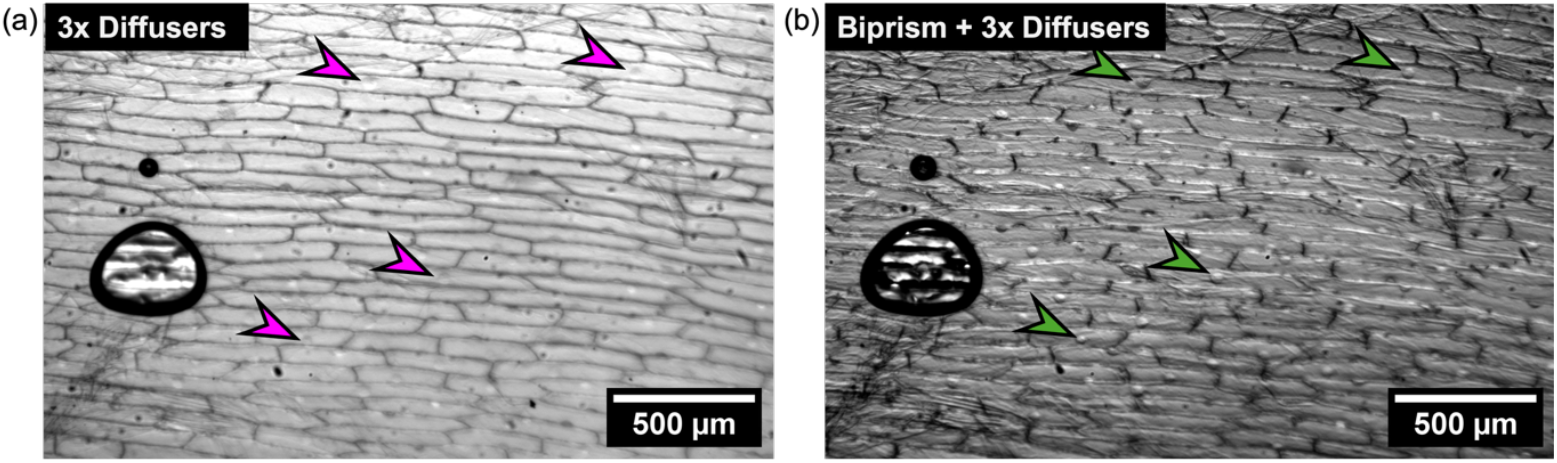
The improved contrast provided by a lens-biprism element introduced into the illumination path of a stereomicroscope reveals biological structure in transparent specimens. **(a)** An image of an onion abaxial epidermis preparation acquired using a transillumination setup with three layered ground glass diffusers. Magenta arrows indicated vaguely discernible cell nuclei. **(b)** An image of an onion abaxial epidermis preparation acquired using a 3D printed biprism transillumination setup. The same nuclei indicated by the green arrows are revealed due to the increased contrast.

## Discussion

We present a 3D printed integrated lens-biprism element which, when placed in the illumination path of a stereomicroscope, provides homogeneous illumination at the specimen plane and facilitates up to a 67.62% contrast enhancement. These improvements permit the visualisation of features otherwise unobserved by conventional stereomicroscopy in optically transparent specimens. Our simple and low-cost modification to the illumination path provides users with a means to visualise refractive features in transparent live specimens across spatial scales.

We observed that the field illumination profile of the standard stereomicroscope was inhomogeneous, with a distinct intensity maximum at the specimen plane that resulted in two laterally offset beams transmitted through the left and right detection axes. While a partial cure was to add diffusive elements, the long-established microscope manufacturers have adopted expensive solutions; for example, having two separate diascopic illuminators of equal intensity, with one for each optical path in the stereomicroscope. Our devised solution uses a 3D printed lens-biprism element to correct the illumination at the specimen plane, which can be easily fabricated at low cost using consumer-grade 3D printers. The materials cost for the 3D printed biprism element amounted to £1.65, with an additional £1.15 for the 3D printed spin chucks. The total cost of the 3D printer and wash/cure station for lens production amounted to approximately £250, with an additional £1200 for the spin coater and vacuum chamber for post-processing the lenses. The cost of the FFF-type printer used to fabricate the spin chucks was approximately £650, although lower-cost options are available.

As expected, we achieved improved contrast due to the improved illumination homogeneity and collimated double side illumination facilitated by the biprism transilluminator compared with that of a diffuser setup. This result resembles that achieved by closing the condenser iris of a standard research-grade non- stereomicroscope (e.g., a brightfield transmission or phase contrast microscope) and, as with such microscopes, makes it easier to find the focus and detect refractive structures^19^.

There are, of course, digital post-processing methods which may improve image contrast, such as Contrast Limited Adaptive Histogram Equalisation (CLAHE)^20^ or AI- driven plugins which inherently increase contrast via denoising (e.g., Noise2Void^21^). However, depending on the specimen, these approaches may lead to the introduction of image processing artefacts, loss of valuable data, or the unwanted amplification of the background signal. Our simple adjustment to the microscope provides means of improving a widely utilised biological imaging system to improve image quality at the point of acquisition, opposed to employing corrective post-processing methods. Ultimately our method minimises the need for corrective post-processing, and any artefacts therein are mitigated.

Advances in metamaterials have seen applications in improving the performance of the stereomicroscope. Long *et al*. recently demonstrated the use of birefringent single-layer metalenses to increase the numerical aperture (NA) and imaging performance of routine stereomicroscope setups^10^. They present a metamaterial objective lens measuring 400 μm in diameter with an NA of 0.4, providing a lateral resolution of 870 nm and an axial resolution of 4 μm. These metalenses provided homogeneous illumination across a limited FoV, but did not correct the illumination over the full field. They were also limited to monochromatic illumination to avoid chromatic aberrations that broadly limit metalens applications so far. It should be noted that metalens fabrication via deposition of thin silicon layers and etching via electron beam lithography is a specialised fabrication method outside of the remit of many stereomicroscope end users and requires expensive instrumentation^22^.

Although this study focused on brightfield transmission imaging of transparent specimens, the lens-biprism element design is inherently compatible with other contrast methods in stereomicroscopy. In polarisation-based contrast methods, such as birefringence imaging or polarised light stereomicroscopy, the lens-biprism element could be positioned prior to the polariser in the illumination path, ensuring that the angularly directed beams remain polarisation-uniform before entering the specimen. The low-birefringence nature of the cured photopolymer resin used in fabrication further supports this compatibility^12,14^. Future adaptations could involve integrating polarising films onto the planar faces of the biprism to extend functionality without altering the detection optics.

The implementation of a low-cost 3D printed lens-biprism element into the illumination path provides enhanced contrast with the routine stereomicroscope. We propose that this simple modification to a commonplace piece of laboratory equipment would serve to increase its value for bioimaging by more reliably discerning structure in transparent specimens. These improvements would not only be valued by the biologist, but also by any discipline that makes use of diascopic stereo-visualisation, including in field-based healthcare diagnostics. Beyond contrast enhancement in transparent biological specimens, the lens-biprism element approach has potential utility in diverse application domains where stereomicroscopes are employed for inspection, assembly, or quality control. In micromanufacturing and microelectronics, for example, improved transmitted-light contrast could assist in detecting microcracks, delamination, or surface defects in transparent or semi-transparent films and substrates. Similarly, in material sciences, the technique could aid in the visualisation of internal stress patterns, anisotropies, or birefringent domains within polymer composites or crystalline specimens when combined with polarised light imaging. For forensic sciences and geological applications, where stereomicroscopy is routinely used to inspect hair fibres, paint flakes, glass fragments, minerals, or tool marks, the biprism- enhanced transillumination setup may improve the detection of low-contrast boundaries or internal structures without requiring staining or destructive preparation. Furthermore, the simplicity and low manufacturing cost makes our biprism transillumination method attractive for use in portable stereomicroscopy systems for applications in the field, especially where illumination quality is often a limiting factor. These possibilities highlight the versatility of the biprism design as a drop-in enhancement to existing optical workflows in both research and applied contexts. Moreover, the gains in image quality afforded by homogeneous illumination may benefit beyond the research lab and into clinical settings, where the low-cost and accessible methods we provide can be readily adopted and implemented.

## Acknowledgements

The Authors wish to thank the funders who supported this work. LMR and GM were funded by the Leverhulme Trust. SF, GWG and GM were funded by the BBSRC BB/X005178/1. JC, CB and RB were funded by BBSRC (BB/Z51486X/1), an EPSRC studentship (EP/T517938/1) and supported by the National Manufacturing Institute Scotland (NMIS). GM was supported by the MRC (MR/K015583/1) and the BBSRC (BB/P02565X/1 and BBT011602).

Figure 1c was created in BioRender. Rooney, L. (2025) BioRender.com/yy1vgqk

## Data Availability Statement

The data that support the findings of this study are openly available at the University of Strathclyde KnowledgeBase.

## Declarations of Interest

The authors declare no conflicts of interest.

## Notes

### Competing Interest Statement

The authors have declared no competing interest.

